# From friend to foe and back - Coevolutionary transitions in the mutualism-antagonism continuum

**DOI:** 10.1101/2024.09.27.615354

**Authors:** Felix Jäger, Frank M. Schurr, Korinna Theresa Allhoff

## Abstract

Interspecific interactions evolve along a continuum ranging from mutualism to antagonism. Evolutionary theory so far focused mostly on parts of this continuum, notably on mechanisms that enable and stabilise mutualism. These mechanisms often involve partner discrimination ensuring that interaction intensity is higher with more cooperative partners. However, the gradual trajectory of coevolutionary transitions between mutualism and antagonism remains unclear. Here, we model how discrimination ability in one partner coevolves with mutualistic service provided by the other and analyse the resulting evolutionary trajectories in the mutualism-antagonism continuum. We show that strong ecological change, such as a radical host shift or colonisation of a new environment, can trigger transitions in both directions including back-and-forth transitions between antagonism and mutualism. Moreover, we find an evolutionary tipping point: a stable mutualism may break down to antagonism if the cost of either mutualistic service or discrimination ability gradually increases above a threshold beyond which this transition cannot be reversed by reducing costs again. Our study provides a new perspective on the evolution of biotic interactions and hence on the dynamic structure of ecological networks.

## 1 Introduction

Interspecific interactions play a key role for ecological and evolutionary dynamics. They can be classified into different interaction types, according to the effect on the fitness of the two involved partners. For instance, antagonism exists in host-parasite relationships, whereas plant-pollinator interactions are an important example for mutualism. However, mutualism and antagonism are not unbridgeable opposites but rather span a continuum with any gradation possible [1, 2]. An interaction’s location in this continuum is by no means fixed. Apart from environmental variation and intraspecific heterogeneity, also evolution changes the costs and benefits for both interaction partners and thereby moves the interaction across the continuum [3, 4]. Eventually, this can result in evolutionary transitions from mutualism to antagonism or vice versa [1, 5].

For example, lineages of originally mutualistic yucca moths evolved into antagonists that stopped pollinating yucca plants but continued ovipositing in the ovary of the flower where the larvae feed on developing seeds [6, 7]. Vice versa, there is compelling evidence of microbial parasites evolving to reduce their virulence, ultimately becoming mutualists [5, 8, 9]. Such shifts of interaction type can have major consequences for community dynamics, ecosystem services or human health [5, 10, 11], and are hypothesised to underlie major evolutionary innovations such as the evolution of pollination [12, 13]. However, a comprehensive theoretical understanding of the gradual evolution of interaction types is still missing [8, 10, 14].

Theoretical work has looked at evolutionary transitions of interaction type mainly through the lens of the classic question: ‘How can mutualism evolve and how can it be maintained in the face of cheaters?’ [15, 16, 17, 18, 19, 20]. While several mechanisms have been identified that potentially enable the evolution of mutualism from antagonism [7, 21, 22, 23], the specific coevolutionary trajectory in the mutualism-antagonism continuum has received little attention. Many models in this context, in particular game-theoretical ones, interpret cooperation and defection as two discrete strategies, ignoring the continuum between them [22, 24, 25, 26, 27, 28]. Studies that instead model the evolution of quantitative traits influencing the interaction type usually do not quantify the precise border between antagonism and mutualism, making it hard to derive conclusions about the course of the transition [29, 30, 31; but see 32].

The focus on the evolution of mutualism introduces a bias in the literature with respect to the direction of transition: very few studies examine the evolutionary shift from mutualism to antagonism [10, 33; but see 34]. Although some empirical results suggest that this occurs less frequently than the transition from antagonism to mutualism [5, 8, 35], inclusion of both transition directions is needed for a comprehensive understanding of dynamics in the mutualism antagonism continuum, also accounting for reversibility of transitions. This holds especially in times of rapid environmental change which triggers diverse coevolutionary adaptations by altering the costs and benefits of an interaction. The resulting potential for mutualism breakdown is a threat for numerous ecosystem services [10].

Here, we present a general model of gradual evolutionary transitions from antagonism to mutualism and vice versa. We track the location of an interaction in the continuum through time. The structure of our model reflects the asymmetry often observed in empirical systems: One partner can potentially transition between antagonist and mutualist by adapting the amount of mutualistic service it provides while the other is able to evolve some discrimination ability, that is, the ability to adapt the strength of interaction according to the service provided by the partner [36, 37]. This comprises different mechanisms thought to promote the evolutionary emergence of mutualism. For instance, partner choice, where one partner can choose beforehand with whom it interacts [15, 19, 38], can be interpreted as conditional adaptation of interaction probability or frequency. Sanctions, where one partner may punish bad-quality partners after engaging in the interaction [39, 40], correspond to a response of interaction duration or intensity to varying cooperativeness.

We use the model to identify pathways of gradual evolutionary transitions of interaction type. In particular, we first analyse the transient coevolutionary dynamics with different initial trait values and examine the effect of varying relative speed of evolution, known to significantly influence any coevolutionary process [41, 42, 43, 44]. In a second step, we then study how varying environmental conditions in the form of varying cost and benefit parameters affect the possibility and course of transitions. We find potential for transitions in both directions as well as back-and-forth transitions, mainly dependent on preadaptation in discrimination ability. Moreover, we observe an evolutionary tipping point: When costs of mutualism or discrimination ability gradually exceed a threshold due to environmental change, a stable mutualism abruptly evolves towards antagonism, with limited reversibility. Our model thus yields qualitatively new results on coevolutionary dynamics in the mutualism-antagonism continuum.

## 2 Model and Methods

### 2.1 Net fitness effects of an interaction

We model a system consisting of two interacting species called ‘host’ and ‘partner’. While the fitness effect of the interaction for the partner is always positive, the host may either benefit or suffer from the interaction, representing mutualism or antagonism, respectively. By explicitly quantifying the fitness effect of the interaction for both host and partner, we are able to track the position of the interaction in the mutualism-antagonism continuum. This includes the special case of commensalism, where the interaction has no effect on host fitness. We assume that the location of the interaction in the continuum is solely determined by traits of the two involved species. Thus, we restrict ourselves to investigate transitions of interaction type resulting from evolutionary adaptation as opposed to transitions due to changing exogeneous biotic or abiotic conditions in context-dependent interactions.

As an example, throughout this section we refer to the nutrient exchange symbiosis between a legume host and a rhizobium. Rhizobia are soil bacteria that infect the roots of legumes where they fix nitrogen (N) from the atmosphere and release it to the plant, while receiving carbon in return. Depending on the level of N they fix, rhizobia can range from mutualists to antagonists [45]. Note that the model equally applies to a variety of other biotic interactions such as between yucca plants and yucca moths (which range from effective pollinators to seed predators; [6]) or between ‘client’ fish and their cleaners (some of which consume host tissue; [46]).

We model the interaction as the reciprocal exchange of services. The partner provides a fitness benefit *b*_*H*_ *≥* 0 to the host (e.g. amount of nitrogen provided by a rhizobium), while the host provides a fitness benefit *b*_*P*_ *≥* 0 to the partner (carbon provisioning by the legume). Both comes at a cost for the respective service provider, which we assume to scale proportionally with the provided service with cost factors *c*_*P*_ *≥* 0 and *c*_*H*_ *≥* 0, respectively. The principal assumption that providing a mutualistic service is costly poses a fundamental conflict of interest between the two species. Although empirical evidence is not unambiguous [16, 47], this is likely to hold in many systems and is the basis for the majority of theoretical research in this context to date [7, 22, 28, 30, 36]. With this simple model we get the following equations for the net fitness effects of one interaction on the host 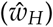 and its partner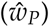, that is, the increase or decrease in per-capita fitness due to one realized interaction:

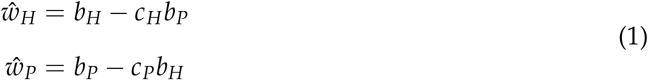

In this study, we focus on the evolution of the service provided to the host by the partner *b*_*H*_. The benefit to the partner provided by the host *b*_*P*_ instead, as well as the cost factors *c*_*H*_ and *c*_*P*_, remain constant throughout the simulations. For simplicity of presentation, instead of *b*_*H*_ we consider the shifted variable *β* = *b*_*H*_ *− c*_*H*_*b*_*P*_, which is the net effect of an interaction on host fitness (e.g. the net effect of a rhizobium on plant fitness). *β* has the property that the interaction can be regarded as beneficial for the host if *β >* 0 and detrimental if *β <* 0. Note that since *c*_*H*_*b*_*P*_ is a constant, *β* can still be interpreted as a partner trait. In the following, we consequently refer to *β* as ‘partner effect’. We get:

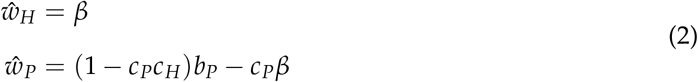

In all subsequent analyses and simulations, we have 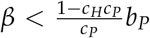 ensuring that the interaction is always beneficial for the partner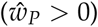.

### 2.2 Host discrimination ability

We extend the presented equations by accounting for discrimination by the host. This is the mechanism that provides potential selection for investment in mutualism by the partner in our model. The host has a trait *α ≥* 0 representing its discrimination ability. *α* measures the extent to which the interaction frequency depends on the net effect of an interaction on host fitness *β* (see Fig. 1). If *α* = 0 (no discrimination ability), interaction frequency is independent of whether the partner is beneficial or detrimental for the host. If *α* is instead large, the interaction frequency *f* (*α, β*) highly depends on the partner trait: It is close to 0 in case of an antagonistic partner (representing avoidance of the interaction) and approximates 1 for a highly mutualistic partner. This relation is mathematically expressed using a sigmoid function:

**Figure 1.**
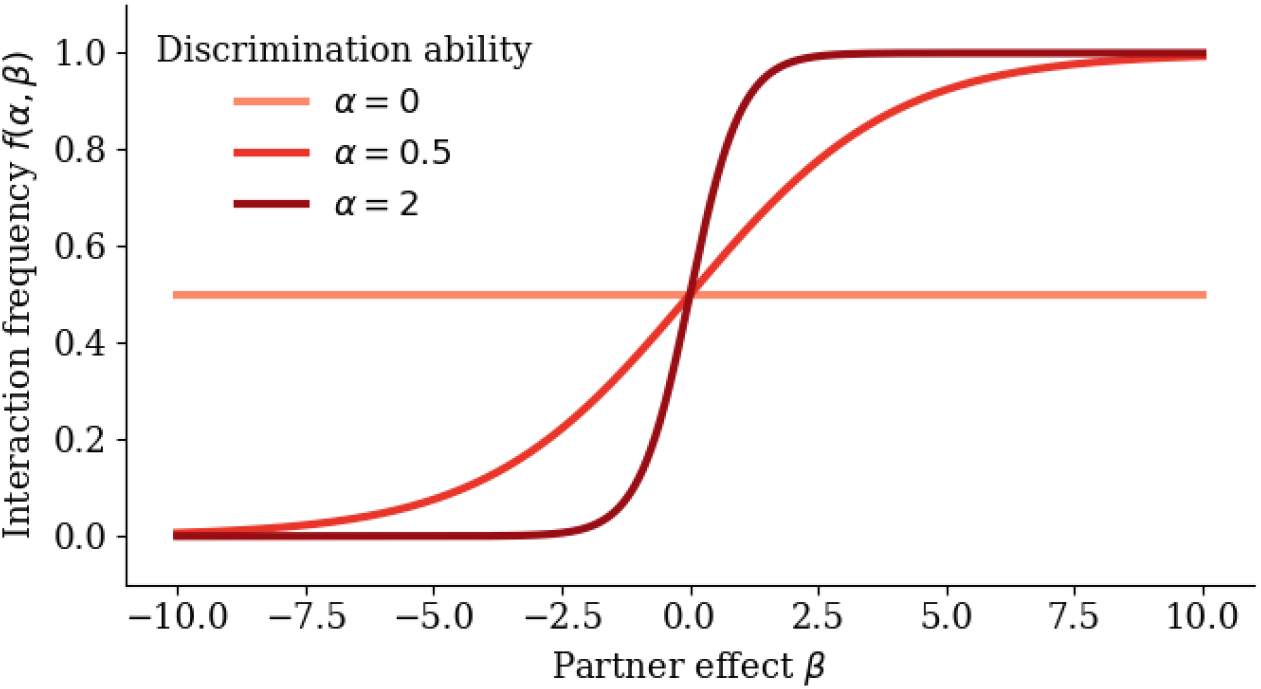
Dependence of interaction frequency on host discrimination ability and partner effect. For small discrimination ability *α*, the influence of the partner effect *β* on the interaction frequency is negligible. For higher *α*, interaction frequency is high with mutualistic partners, but low with antagonistic ones.

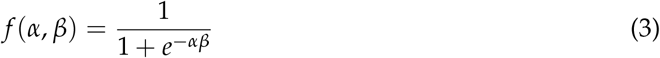

Note that *f* (*α, β*) can alternatively be interpreted not as interaction frequency, but interaction intensity or duration which broadens the range of biological mechanisms that are potentially captured by our concept of discrimination.

In nature, discrimination ability *α* represents the accuracy with which the plants direct carbon to mutualistic as opposed to non-mutualistic rhizobia [48] (or mycorrhiza; [49]). Other examples include the accuracy, with which corals provide nitrogen to cooperative vs. selfish endosymbionts [50], with which yucca plants retain fruits containing yucca moths that are effective pollinators rather than seed predators [51], or with which ‘client’ fish tolerate mutualistic cleaner fish (that remove ectoparasites) as opposed to cheaters (that consume host tissue; [46]).

The final equations describing the net fitness effects on host (*w*_*H*_) and partner (*w*_*P*_), while accounting for varying interaction frequency, are the following. Note that we decided to use a model without explicit population dynamics (see Supplementary Material S.1 for justification).

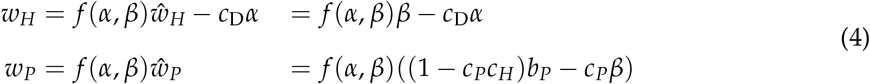

They arise from equations 2 by multiplying the fitness effects of one realized interaction (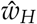 and 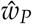) with the interaction frequency *f* (*α, β*), which depends on both host discrimination ability *α* and partner phenotype *β* as outlined above. Additionally, host discrimination ability comes at a cost for the host that is proportional to *α* with cost factor *c*_*D*_ (cost of discrimination ability). For example, this could be an energetic cost for processing the signal about the N fixation level in the molecular dialogue between legume and rhizobium.

### 2.3 Evolutionary dynamics

The simultaneous changes in mean trait value of both host discrimination ability *α* and partner effect *β* are described by differential equations:

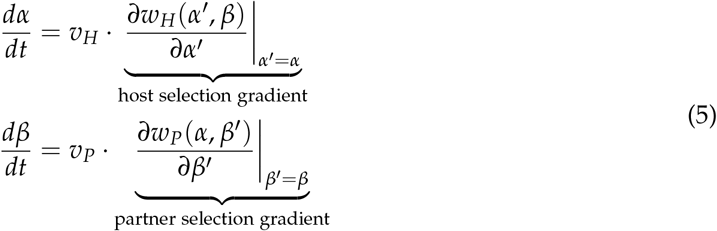

These equations arise from quantitative genetics under the assumption of small trait variance [52, 53, 54] and resemble the Canonical Equation of adaptive dynamics [55]. Trait change is proportional to the selection gradient, the local gradient of the fitness function with respect to the focal trait. *v*_*H*_ and *v*_*P*_ are the additive genetic variances for the host and partner trait divided by the respective generation time [43]. They represent the speed of evolutionary adaptation in host and partner, assumed to be constant over time, and influenced by rate and effect size of mutations. To prevent biologically implausible negative values of discrimination ability *α*, we set 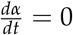 in case of *α* = 0 and a negative host selection gradient.

Note that we assume that only the per-capita fitness effects of the interaction *w*_*H*_ and *w*_*P*_ determine selection in our model. We thereby neglect the influence of other selection pressures on the focal traits as well as potential density-dependent selection (see Supplementary Material S.1 for alternative model formulations).

### 2.4 Model analysis

First, we analyse the evolutionary dynamics in the model using the baseline parameterisation given in Table 1. We start by examining the sign of the selection gradients for host and partner (see equations 5) depending on host discrimination ability *α* and partner effect *β*. In particular, we focus on the shape of the evolutionary null-isoclines where the host or partner selection gradient equals zero. The intersections of host and partner null-isocline represent evolutionary fixed points, which are potential endpoints of evolution since selection pressures vanish for both species. Using the eigenvalues of the Jacobian of the system described by equations 5, we assess the local asymptotic stability of the fixed points, i.e. whether evolution will let a system close to a fixed point converge to it or not.

**Table 1:** Model parameters and their standard values.

| Parameter | Meaning | Standard value |
| --- | --- | --- |
| $b_P$ | benefit for the partner provided by the host | 5 |
| $c_H$ | cost of mutualistic service for the host | 0.6 |
| $c_P$ | cost of mutualistic service for the partner | 0.6 |
| $c_D$ | cost of discrimination ability for the host | 0.1 |
| $v_H$ | speed of host adaptation | 0.02 |
| $v_P$ | speed of partner adaptation | 0.02 |

The insights gained into the system’s behaviour subsequently enable us to understand the evolutionary transitions from antagonism to mutualism and vice versa in the transient dynamics of the model (i.e. when starting far from an evolutionary equilibrium). We compare the potential for evolutionary transitions in the transient dynamics for varying relative speed of evolution, mediated by a varying ratio of *v*_*H*_ and *v*_*P*_.

Finally, we aim at identifying potential evolutionary transitions resulting from gradual change in the biotic and abiotic environment. We interpret environmental change implicitly as a modification of the cost and benefit parameters *c*_*H*_, *c*_*P*_, *c*_*D*_ and *b*_*P*_. For instance, in the legume-rhizobium interaction, a long-term nitrogen enrichment of the soil would correspond to an increase of costs for the rhizobium for providing the same fitness benefit to the plant, since the plant can use soil N more easily.

The outlined analyses already account for variation of any model parameter which ensures the generality of the obtained results (see Supplementary Material S.1).

### 2.5 Individual-based simulations

We complement the outlined model by individual-based simulations that relax some of the strict assumptions of the analytical approach (see Supplementary Material S.2). In particular, we introduce demographic stochasticity and intraspecific variation, as well as the possibility of large-effect mutations to the system. Each of these aspects can potentially alter the evolutionary dynamics fundamentally [41, 56, 57, 58]. Notably, the individual-based model allows for comparison of scenarios with varying effect size of mutations. The implementation is closely oriented to the analytical model in order to ensure comparability.

## 3 Results

### 3.1 Two evolutionary attractors

We first examine the sign of the two selection gradients and the shape of the evolutionary null-isoclines. This enables us to understand the selection mechanisms that underlie potential transitions in the mutualism-antagonism continuum in the model. Figure 2a shows the sign of the selection gradient for the partner trait, the net effect of an interaction on host fitness *β*, depending on host discrimination ability *α* and *β*. A positive selection gradient means selection for increased mutualism, a negative selection gradient represents selection for more antagonistic values of *β*. Without any discrimination by the host (*α* = 0), selection is always negative due to the cost of mutualism for the partner. However, with increased discrimination ability, the partner experiences selection for higher values of *β* in order to prevent being rejected by the host. The border between the region of positive and negative selection is the evolutionary null-isocline for the partner. Note that for very high values of discrimination ability *α* it suffices for the partner to be only slightly mutualistic to retain full interaction frequency by the host (see Fig. 1). Therefore, since we assume that partners providing more benefits for the host face greater costs, the partner null-isocline approximates commensalism (*β* = 0).

**Figure 2.**
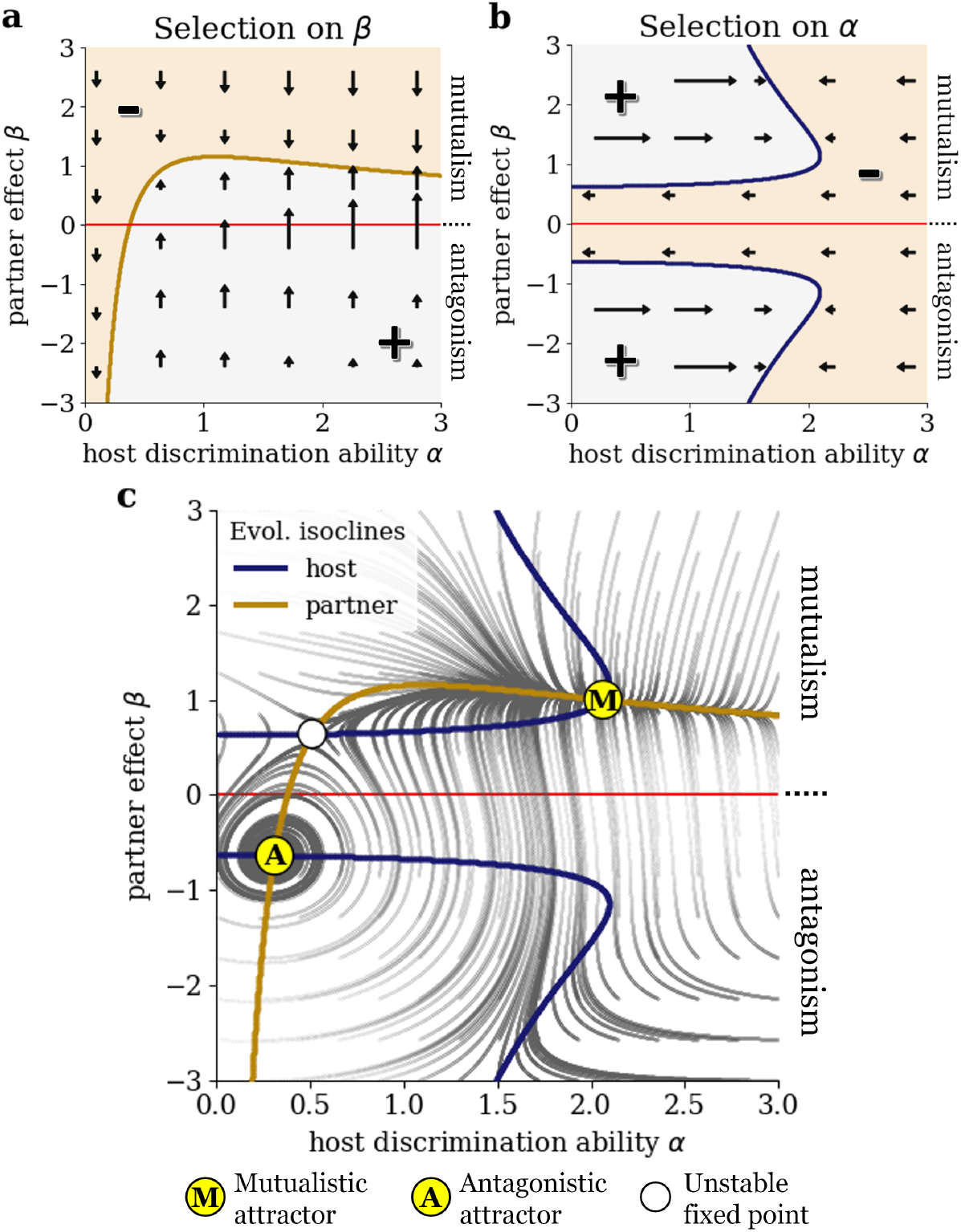
Coevolution of partner effect and host discrimination ability along the mutualismantagonism continuum. Colour plots show the sign of the selection gradients for (**a**) the net effect of an interaction on host fitness *β* and (**b**) the host discrimination ability *α*, depending on *α* and *β* (all other parameters are chosen according to Table 1). Positive selection gradient is shown in light grey, light brown colour indicates negative selection gradient. Arrows visualise the direction and strength of selection. The blue and brown line represent host and partner evolutionary null-isocline, respectively, where selection on *α* or *β* vanishes. (**c**) The intersections of the null-isoclines are the stable/unstable evolutionary fixed points, marked by yellow/white circles. The dark grey lines visualise the evolutionary trajectories for varying initial values of discrimination ability *α* and partner effect *β*.

Next, we look at the selection gradient for the discrimination ability of the host *α* and the corresponding evolutionary host null-isocline (Fig. 2b). For *β* = 0, the interaction does not affect host fitness, hence there is no need to invest in discrimination: Selection is always negative. However, if the partner effect *β* is positive or negative enough, a certain degree of discrimination becomes worthwhile for the host. This either reflects selection for increasing interaction frequency with a mutualist (region of positive selection above x-axis) or selection for avoidance of an antagonist (region of positive selection below x-axis). For very high or very low values of *β*, already a slight investment in discrimination ability renders the discrimination almost maximally efficient, as can be seen from the shape of the interaction frequency function *f* (see Fig. 1). Therefore, the host null-isocline approaches the y-axis (no discrimination) asymptotically.

We find three evolutionary fixed points at the intersections of the null-isoclines in the trait space (the space spanned by the traits *α* and *β*), as shown in Fig. 2c. A linear stability analysis using the eigenvalues of the Jacobian of the system reveals that two of them are locally asymptotically stable (indicated by yellow dots), meaning that evolutionary trajectories starting nearby eventually end up at the fixed point. This is confirmed by numerical simulations of the coevolutionary dynamics for varying initial combinations of host discrimination ability and partner effect, the trajectories of which are plotted in Fig. 2c. One of the two attractors is located in the antagonistic region (*β <* 0) at a low value of *α* while the other is a mutualistic attractor (*β >* 0) with a considerably higher discrimination ability of the host. Additionally, there is an unstable mutualistic fixed point (indicated by a white dot).

The eigenvalues at the antagonistic attractor have a non-zero imaginary part (*−*9.0 *·* 10^*−*4^ + 1.1 *·* 10^*−*2^*i* and *−*9.0 *·* 10^*−*4^ *−* 1.1 *·* 10^*−*2^*i*). This is consistent with the fact that evolutionary trajectories approaching this fixed point show oscillatory behaviour: Increased discrimination ability of the host selects for higher benefits provided by the partner, which in turn selects for lower host discrimination ability since discrimination is not needed when the interaction approaches commensalism. Lack of discrimination selects for antagonistic behaviour of the partner, putting positive selection pressure on discrimination ability again and the loop starts all over. By contrast, the eigenvalues at the mutualistic attractor are real numbers (*−*1.7 *·* 10^*−*3^ and *−*2.2 *·* 10^*−*2^), indicating non-oscillatory trajectories close to this attractor.

### 3.2 Relative speed of evolution governs transitions of interaction type

We closer examine the potential for evolutionary transitions along the mutualism-antagonism continuum in the transient dynamics of the system and compare scenarios with varying relative speed of evolution *v*_*H*_/*v*_*P*_. The adaptation speeds do not change the sign of the selection gradients and consequently have no effect on the evolutionary null-isoclines and the location of evolutionary fixed points in the trait space. However, the ratio *v*_*H*_/*v*_*P*_ alters the shape of the evolutionary trajectories and therefore also the boundary between the basins of attraction (the set of initial conditions for which the system converges to a given attractor). The trajectories for varying relative speed of evolution are shown in Figure 3. Trajectories are coloured blue if they converge to the mutualistic attractor and red if they end up at the antagonist attractor, visualising the basins of attraction. A transition from mutualism to antagonism or vice versa corresponds to a change of sign of *β*, i.e. a crossing of the x-axis.

**Figure 3.**
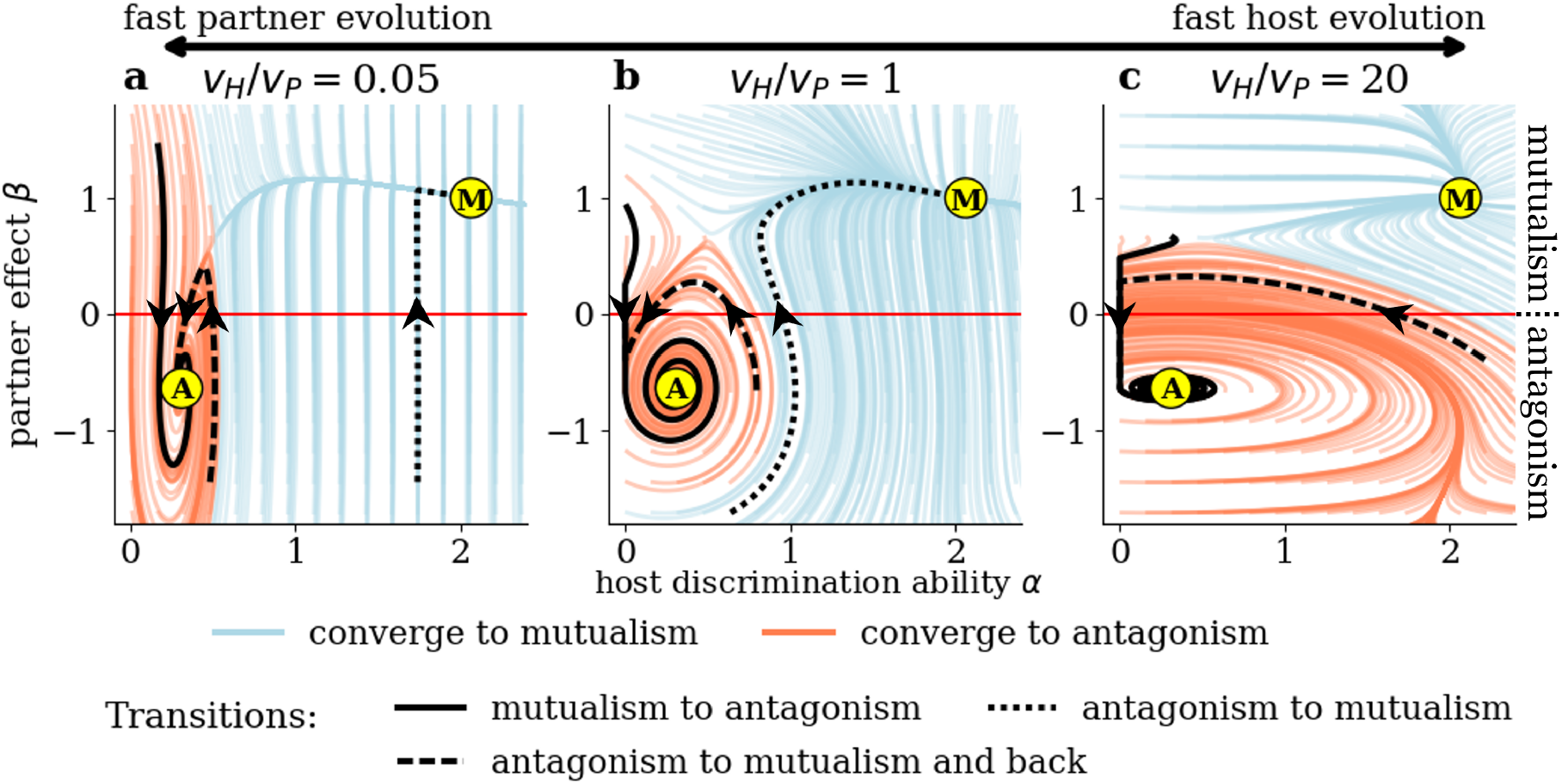
Evolutionary transitions between mutualism and antagonism depend on the relative speed of evolution in hosts and partners. Evolutionary trajectories are shown for (a) fast partner evolution (*v*_*H*_ = 0.001, *v*_*P*_ = 0.02), (b) comparable speed of evolution (*v*_*H*_ = *v*_*P*_ = 0.02) and (c) fast host evolution (*v*_*H*_ = 0.4, *v*_*P*_ = 0.02; all other parameters are chosen according to Table 1). Trajectories are coloured light blue if they end up at the mutualistic attractor (M) and red if they end up at the antagonist one (A), thereby visualising the basins of attraction. Example trajectories highlight the three types of evolutionary transitions of interaction type described in the main text.

In general, there are three different pathways for evolutionary transitions from antagonism to mutualism or vice versa (see highlighted trajectories in Fig. 3):

1. An initially mutualistic partner becomes antagonistic because of the host’s low initial discrimination ability.
2. An initially antagonistic partner becomes mutualistic because of the host’s high initial or evolved discrimination ability.
3. An initially antagonistic partner first becomes mutualistic due to initial or evolved host discrimination ability, but not mutualistic enough to prevent further evolution of reduced discrimination ability; the partner finally becomes antagonistic again (back-and-forth-transition).

For a fast evolving partner (*v*_*H*_/*v*_*P*_ = 0.05, see Fig. 3a), the outcome of the evolution strongly depends on the initial discrimination ability of the host. If *α* is low, the partner will quickly evolve antagonist behaviour and the system ends up at the antagonistic attractor. In particular, high speed of partner evolution increases the occurrence of transitions from mutualism to antagonism. However, if there is enough preadaptation in discrimination ability of the host (*α* ≳ 0.55), there is selection for the partner to exhibit more mutualistic behaviour and trajectories end up at the mutualistic attractor (potential for transitions from antagonism to mutualism). There is only a small window of initial conditions allowing for a back-and-forth transition.

If host and partner evolve at comparable speeds (*v*_*H*_/*v*_*P*_ = 1, Fig. 3b), stable mutualism can evolve even with an initially non-discriminating host. This can happen in two ways. First, if the partner starts as a sufficiently strong mutualist, evolution of the host’s discrimination ability is fast enough to prevent evolution of antagonism in the partner. Secondly, surprisingly, if the partner initially is very antagonistic, strong positive selection lets discrimination ability of the host increase so much that the partner is finally driven towards mutualism. For moderate antagonistic or slightly mutualistic initial values of *β*, evolution still ends up at the antagonistic fixed point (potential for transitions from mutualism to antagonism and back-and-forth transitions).

Finally, if the host evolves much faster than the partner (*v*_*H*_/*v*_*P*_ = 20, Fig. 3c), the basin of attraction of the antagonist attractor expands and covers large parts of the antagonist quadrant, almost ruling out transitions from antagonism to stable mutualism. This is the case because as soon as a partner approaches commensalism (*β ≈* 0), host discrimination ability is selected against and its fast decrease prevents the evolution of stable mutualism.

### 3.3 Changing costs cause evolutionary tipping points

In order to detect evolutionary transitions between mutualism and antagonism triggered by environmental change, we vary the cost parameters *c*_*P*_ and *c*_*D*_ in our model. Figure 4a shows the trait space with evolutionary null-isoclines for different levels of costs of mutualism for the partner *c*_*P*_. Higher cost of mutualism flattens the partner null-isocline and pushes it towards antagonism. This reflects that selection on the service provided by the partner is dominated more and more by its costs instead of its benefits for the partner mediated by discrimination. For the evolutionary fixed points (the intersection of the evolutionary null-isoclines), this implies that for increasing *c*_*P*_ the two fixed points in the mutualistic region approach each other until they culminate in a unique fixed point at around *c*_*P*_ *≈* 0.8, vanishing for higher values of *c*_*P*_ and leaving only the antagonistic fixed point.

**Figure 4.**
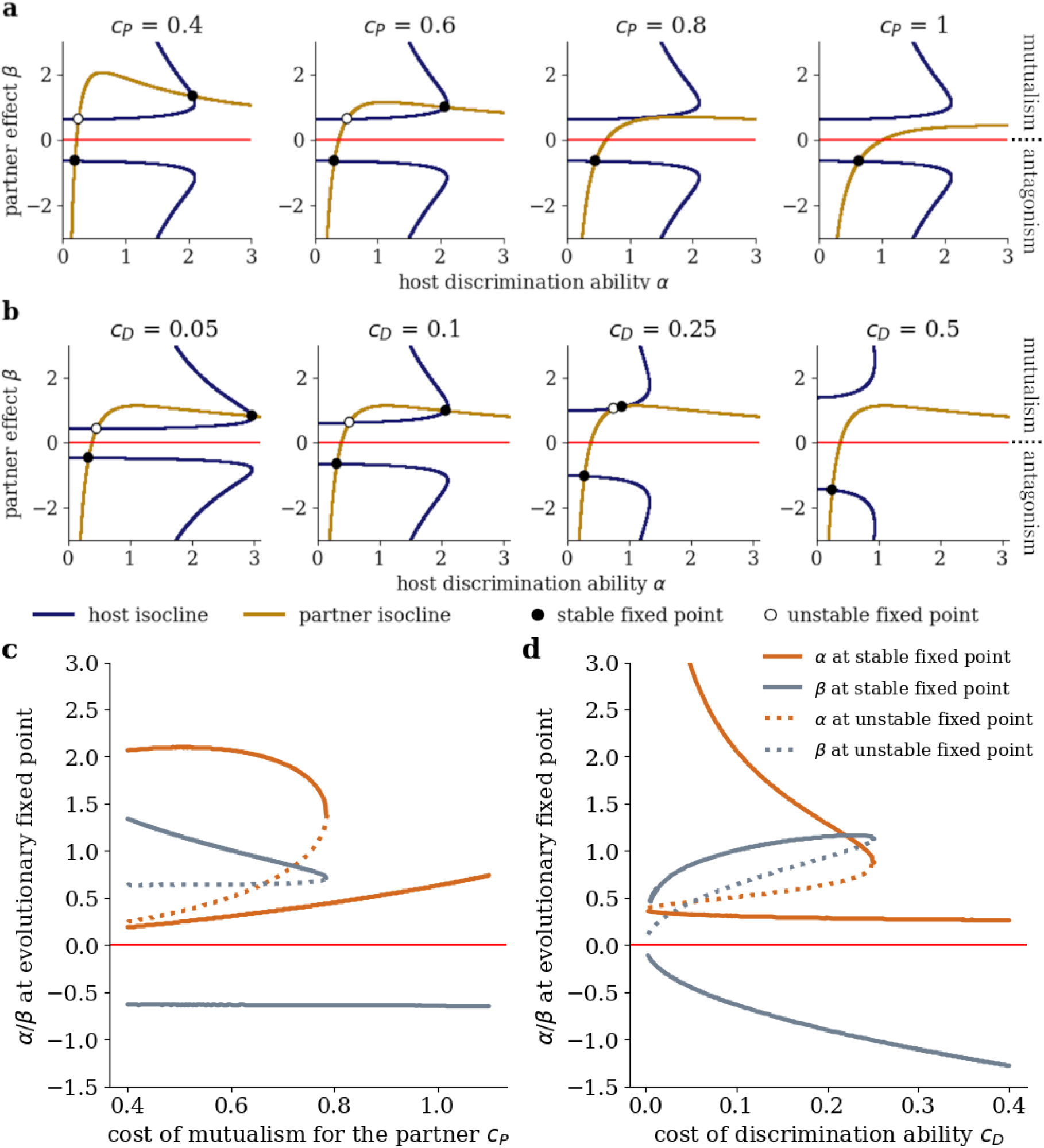
Changes in interaction costs cause evolutionary tipping points. The evolutionary null-isoclines change their shape in the trait space for varying cost of mutualism for the partner *c*_*P*_ (**a**) and varying cost of discrimination ability for the host *c*_*D*_ (**b**). Black/white circles mark the stable/unstable evolutionary fixed points. The baseline scenario corresponds to *c*_*P*_ = 0.6 and *c*_*D*_ = 0.1. Bifurcation diagrams show the trait values at evolutionary fixed points for varying cost of mutualistic service *c*_*P*_ (**c**) and cost of discrimination ability *c*_*D*_ (**d**). Increasing the costs eliminates the mutualistic fixed points and leaves the antagonistic fixed point as the unique attractor.

The described collision of fixed points can be visualised in a bifurcation diagram showing the trait space coordinates of the fixed points depending on the parameter *c*_*P*_ (Fig. 4c). The change in system behaviour we observe at *c*_*P*_ *≈* 0.8 is a saddle-node bifurcation [59] and may represent an evolutionary tipping point [60]: With increasing cost of mutualism for the partner, an interaction which resides at the stable mutualistic attractor is suddenly disrupted and evolves towards the antagonistic fixed point. Crucially, a subsequent reduction of cost cannot reverse the process.

Note that the same evolutionary tipping point is caused by a gradual increase of the cost of mutualism for the host *c*_*H*_ or by a gradual reduction of the benefit *b*_*P*_ for the partner. This is due to the fact that increasing *c*_*H*_ and decreasing *b*_*P*_ have the same effect on the sign of the partner selection gradient as increasing *c*_*P*_, which can be seen from equations 4 (see Supplementary Material S.1). Also note that another evolutionary tipping point arises when costs of discrimination ability for the host *c*_*D*_ increase (Fig. 4b). In this case, the host null-isocline is pushed towards lower discrimination ability *α* and larger absolute values of partner effect *β*. Eventually, the stable and unstable mutualistic fixed point collide and dissolve at *c*_*D*_ *≈* 0.26 (Fig. 4d).

Interestingly, we find that no change of parameter values leads to a gradual transition of a stable fixed point from antagonism to mutualism or vice versa. As a centrepiece of such a transition, a stable commensalistic fixed point would be needed for some parameter combination. This cannot occur in our model since the host null-isocline never crosses the commensalistic axis. In a commensalistic interaction, host discrimination ability *α* is always selected against and if it is reduced to 0, the partner evolves towards antagonism.

### 3.4 Individual-based simulations confirm model predictions

In general, the individual-based simulations corroborated the qualitative results from the analytical model presented in the above sections. However, by introducing stochastic mutations, they enabled trajectories to jump from one basin of attraction to the other, providing an additional source of transitions from antagonism to mutualism or vice versa. The occurrence of such jumps was positively correlated with the effect size of mutations and thus with the amount of intraspecific trait variation present in the model (see Supplementary Material S.3).

## 4 Discussion

The coevolution of discrimination ability of a host and mutualistic service provided by its partner entails gradual evolutionary transitions in the mutualism-antagonism continuum. We identified three different transition pathways under constant environmental conditions in the transient dynamics of the system when starting far from an evolutionary equilibrium state. Moreover, we detected an evolutionary tipping point: A stable mutualism is prone to an abrupt breakdown to antagonism when costs of mutualistic service or costs of discrimination ability gradually increase in a changing environment. The robustness of these results was confirmed by stochastic individual-based simulations. In the following, we first discuss the necessary prerequisites for evolutionary transitions in the transient dynamics before outlining current theoretical developments and potential empirical tests for coevolutionary tipping points.

### 4.1 Determinants of transitions along the mutualism-antagonism continuum

Radical ecological change, such as a switch of the host species, colonisation of a new environment or changes in the community composition, can push biotic interactions far from an evolutionary equilibrium [5, 10]. In the transient dynamics of our model we could observe evolutionary transitions between mutualism and antagonism following such radical change. Notably, the model allowed us to investigate the role of the initial host discrimination ability for such transitions. This is particularly interesting in light of the open question whether preadaptations of host discrimination ability are a necessary prerequisite for the evolution of mutualism [61, 62].

Our results emphasise the importance of preadaptations of host discrimination ability in case of fast-evolving partners. For instance, microbial symbionts most often evolve much faster than their hosts due to smaller generation times [5]. The occurrence of transitions of interaction type then crucially relies on the initial host discrimination ability. If the interaction frequency or duration is not adapted to the amount of service provided by the partner in the first place, which is most likely to be the case in newly assembled interactions [63], an initially mutualistic partner will evolve towards antagonism. Transitions from antagonism to mutualism are possible only if the host already possesses discrimination ability. For example, legumes probably allocated carbon to roots in response to local nitrogen availability before associating with rhizobia in the first place, thereafter selecting for rhizobia to evolve high nitrogen fixation levels [19, 64]. Similarly, the selective abortion of yucca flowers depending on pollen quantity is considered to be a preadaptation rather than a coevolved response to the interaction with yucca moths [19]. However, in systems where host and partner evolve at comparable speeds, e.g. in the interaction between short-lived plants and mutualistic or antagonistic flower visitors [65], host discrimination ability can also build up during the coevolutionary process enabling transitions to mutualism without preadapted discrimination ability.

Interestingly, after a transition from antagonism to mutualism, the interaction does not necessarily stay mutualistic. It approaches a stable mutualistic state only if the initial discrimination ability of the host is high enough. Then the partner invests in mutualism because it otherwise would be discriminated against and the host invests in discrimination ability to maintain high interaction strength with the partner. Otherwise, if the partner does not become sufficiently mutualistic, discrimination is selected against because the host is not strongly affected by the interaction. Consequently, discrimination ability of the host and thereafter also the service provided by the partner reduce and the interaction oscillates back to antagonism. Such back-and-forth transitions along the mutualism-antagonism continuum may be common [5, 30] but hard to detect in empirical research since transitions in both directions need to be verified and since the transient mutualism is weak.

Our results crucially depend on our notion of discrimination ability. This generalises various mechanisms such as partner choice, sanctions and partner fidelity feedback that have been proposed to enable the evolution of mutualism by reducing them to the same underlying principle: The response of the host depends on the service provided by the partner [33, 62]. This basic similarity is presumably the main cause for the striking lack of clear distinction of the mechanisms in the literature [19]. For instance, the selective abortion of flowers by yucca plants was given as an example for each of the three mechanisms mentioned above [22, 23, 36, 66].

However, there is a subtle difference between our model of discrimination and previous conceptions regarding the role of intraspecific variation of partner quality. In contrast to previous models of partner choice, for example, the host does not choose between different present partner individuals. Instead, it adapts the interaction strength with one present partner without any alternatives. By presenting a model where selection for discrimination is not driven by intraspecific trait variation, we provide a potential solution to the often-cited paradox that discrimination ability erases its own selective incentive by eliminating variation in partner quality [7, 20, 30].

We acknowledge that including substantial trait variation in our model could significantly change our results. For instance, if a host is faced with mutualistic and antagonistic partners at the same time, there should be selection for discrimination although the mean partner trait might be close to commensalistic. The presented individual-based model relaxes the assumption of monomorphic populations and suggests that increased trait variation increases the likelihood of additional stochastic transitions of interaction type. Still, overall levels of trait variance remained rather low (see Supplementary Material S.3). A thorough analysis of the influence of trait variation, for example achieved by varying mutation frequency and amplitude, is worth further investigation.

### 4.2 Tipping points in the evolution of biotic interactions

In our model, a gradual change of the environmental parameters does not lead to an equally gradual transition of a stable fixed point from mutualism to antagonism or vice versa. This is because the model never exhibits stable commensalism; commensalism is always a merely transient state. Seemingly contradicting the prevalence of commensalism in nature [67], a potential explanation might be that the weak stable antagonism predicted by our model for low costs of discrimination is empirically indistinguishable from commensalism. More empirical and theoretical research is needed on the stability of commensalism and its significance as a waypoint along the mutualism-antagonism continuum.

Instead of a gradual transition we find an abrupt shift from a stable mutualism to antagonism when costs of mutualistic service for the host or the partner or costs of discrimination ability for the host increase above a certain threshold. Such a transition is characterised by a positive self-reinforcing feedback loop: Under the new environmental conditions, reduced service provisioning by the partner selects for reduced discrimination ability of the host, which in turn selects for more antagonistic behaviour of the partner. Finally, the interaction ends up at an antagonistic equilibrium and a subsequent reduction of costs does not restore the original mutualism. Such hysteresis is inextricably linked to the occurrence of tipping points [68].

Tipping point theory has proven successful in ecology by providing an explanation for abrupt regime shifts in population or community dynamics [69], and the role of evolutionary dynamics in shaping such catastrophic transitions is increasingly recognised [60, 70]. However, only few studies have identified evolutionary tipping points in the sense that a system is driven to an alternative stable state in the trait space (note that the term ‘evolutionary tipping point’ has also been used differently [71]). For example, plant attractiveness in a plant-pollinator interaction was shown to abruptly reduce if pollinator intrinsic growth rate falls below a threshold [72]. The application of tipping point theory to systems of multiple coevolving traits remains to the best of our knowledge unexplored. Ultimately, we believe that evolutionary tipping point theory can help to explain fundamental empirical patterns such as the observation that evolution often happens in short dynamic time intervals (or “punctuated equilibria” [73]) followed by longer steady periods [74, 75, 76].

To date, we are not aware of any empirical attempt to detect a coevolutionary tipping point. We see two possibilities to test the predictions of our model. The first approach is based on controlled coevolution experiments with gradual variation of environmental conditions and quantification of the fitness effect of the interaction for the host in order to determine the interaction type. Some studies using the legume-rhizobium model system showed that increased nitrogen availability in the soil can lead to rapid evolution of reduced cooperativeness of rhizobia [77, 78, 79], potentially reflecting the dynamics found in our model. Here, a nitrogen-rich environment corresponds to higher costs of mutualistic service for the rhizobium since it needs to provide more nitrogen to achieve the same fitness benefit for the host plant, given that more alternative resources are available. However, no gradient of different soil treatments was used in these experiments and no test for reversibility was carried out, needed to unequivocally identify a tipping point. Alternatively, the existence of the tipping point might also be inferred from macroevolutionary data on changes in interaction type between periods that gradually differed in interaction costs. Ideally, data sets comprise periods with environmental change in both directions to analyse reversibility of the transitions of interaction type.

A major challenge in both described approaches would be to disentangle the influence of evolution (in response to environmental change) and the context-dependency of the interaction type, present without any evolution at play. Furthermore, the question remains whether evolution of costs (which are assumed fixed in our model) would push real-world system towards the described tipping point. One might reasonably assume that evolution leads to reduced costs. For example, rhizobia might evolve to fix the same amount of nitrogen at a lower metabolic cost to themselves or plants evolve reduced costs of nectar production. These reduced costs could then even enable evolutionary transitions from antagonism to mutualism. Our model suggests that for reduced costs, the antagonistic stable fixed point persists, but decreases in stability, meaning that stochastic transitions towards mutualism become more and more likely in the presence of intraspecific variability (see Fig. 4). In a similar fashion, the trajectories in the mutualism-antagonism-continuum could also be affected by the evolution of increased benefit to the partner provided by the host. Future extensions of our work should thus account for the evolution of all the involved costs and benefits, focussing in particular on the role of intraspecific variability.

## 5 Conclusion

Mutualism and antagonism are increasingly recognised as spanning a continuum of possible outcomes of an interspecific interaction. We developed a theoretical framework to study how coevolution moves a two-species interaction along this continuum. We found potential for transitions from mutualism to antagonism and vice versa, depending on the initial discrimination ability of the host or in the form of evolutionary tipping points when costs gradually change. Our study thereby formulates general hypotheses for empirical work on shifts of interaction type in diverse contexts ranging from pollination to nutrient exchange symbioses. Since the interaction type has significant influence on structure and stability of complex ecological networks [80, 81], our results should have important implications for the study of communities consisting of multiple mutualistic and antagonistic interactions [11, 82, 83].

