## Supplementary Material for "From friend to foe and back - Coevolutionary transitions in the mutualism-antagonism continuum"

### S.1 Generality of model results using alternative model setups

In this section, we give justification for different model assumptions. We first explain why we refrain from modelling explicit population dynamics. Then we outline why, within the chosen model, no further variation of parameters is needed to ensure the generality of the obtained results.

As an alternative to the outlined setup, other model versions were tested using explicit population dynamics. These include intrinsic growth and competition processes of the two populations. Change of host density $H$ and partner density $P$ were described by the following Lotka-Volterra-type differential equations:

$$\frac{dH}{dt} = H(r_H - a_H H + w_H P)$$
$$\frac{dP}{dt} = P(r_P - a_P P + w_P H),$$

where $r_H$ and $r_P$ are intrinsic growth rates of host and partner, $a_H$ and $a_P$ coefficients for intraspecific competition within host and partner populations. Using the framework of adaptive dynamics [1, 2, 3], from such explicit population dynamics it is possible to derive a canonical equation for evolutionary dynamics similar to equations 5 from the main text, accounting for potential density-dependent selection [4]. In our case, the main difference is that selection gradients are additionally multiplied with equilibrium densities of the respective species. However, this does not change the sign of the selection gradient and hence not the evolutionary null-isoclines and fixed points described in the main text. Differences with respect to the transient dynamics were marginal, hence, for the sake of simplicity, we decided to use a model without explicit population dynamics.

The results presented in the main text are generic in the sense that they account for any effects that can occur due to variation of any model parameter. Varying the values of the cost parameters $c_D$ (cost of discrimination ability for the host) and $c_P$ (cost of mutualistic service for the partner), as well as varying the adaptation speeds $v_H$ and $v_P$ was part of the analysis of the model presented in the main text. From equations 4 from the main text one can see that a variation of the partner benefit $b_P$ or the cost of mutualistic service for the host $c_H$ corresponds to a variation of $c_P$ combined with rescaling $w_P$ by a constant factor. Also, changing the maximal interaction frequency in the function $f(\alpha, \beta)$ would result in a proportional rescaling of $w_H$ and $w_P$. Rescaling does not change the sign of the selection gradient; moreover, when looking at the trait change dynamics described by equations 5 from the main text, it simply corresponds to a change of adaptation speeds $v_H$ and/or $v_P$. This was investigated in the main text.

### S.2 Implementation of the individual-based simulations

We built an individual-based model (IBM) to complement the analytical model by introducing demographic stochasticity, intraspecific variation and large-effect mutations. Typically, IBMs used in similar contexts make birth and death rates explicit, often based on the Gillespie algorithm [5, 6, 7]. Here, we neglect instead the population dynamics and apply fixed population sizes in order to ensure comparability with the analytical approach. The model was implemented in NetLogo 6.4.0 and is publicly accessible [8].

The IBM comprises a population of $N = 100$ hosts and $N = 100$ partners. The population sizes remain constant throughout the simulations. Host individuals are characterised by their discrimination ability $\alpha$, partners by their net effect on host fitness $\beta$. The model is initialised with monomorphic populations (meaning that all individuals share the same trait value), but experiences intraspecific diversification during the simulations. Each time step consists of an interaction phase and an asexual reproduction event.

- **Interaction phase.** Hosts and partners are divided into 100 host-partner pairs that are assembled at random. For each pair, the interaction is considered successful with probability $f(\alpha, \beta)$, calculated based on the trait values $\alpha$ and $\beta$ of the two involved individuals using equation 3 from the main text. In case of a successful interaction, host "fitness" increases by $\beta - c_D\alpha$ and partner "fitness" increases by $(1 - c_P c_H)b_P - c_P\beta$ ("fitness" here is a performance measure without unit, later used to determine which individual is more likely to reproduce). If the interaction is not successful, host fitness decreases by $c_D\alpha$ whereas partner fitness remains unchanged. This reflects the analytical model (equation 4 from the main text) where cost of discrimination is independent of interaction frequency. The process is repeated five times within one timestep with varying subdivisions into pairs. The total fitness (the added fitness from the five rounds of interaction) is rescaled such that the minimum total fitness is 0 in both populations.

- **Reproduction event.** In even timesteps, a host individual reproduces. One parent host is drawn from the host population, with probability proportional to the total fitness from the interaction phase. An offspring individual is created with discrimination ability $\alpha$ drawn from a normal distribution around the corresponding trait value of the parent; the standard deviation is given by the parameter $\sigma_\alpha$. Potential negative values of $\alpha$ are replaced by 0. Another host individual dies at random.
  Partners reproduce in exactly the same way but in odd timesteps (the standard deviation of mutation distribution being $\sigma_\beta$, and with negative $\beta$-values allowed). After reproduction, host and partner fitnesses are reset to 0.

The parameter values in the baseline scenario were the same as for the analytical model (see Table 1 from the main text), the default value for $\sigma_\alpha$ and $\sigma_\beta$ was 0.05. By tracking the mean discrimination ability $\alpha$ and mean partner effect $\beta$ over time, we were able to investigate the emerging evolutionary dynamics in the IBM. These were then compared to the results of the analytical model. In addition to varying the parameters from the analytical model, we analysed the influence of mutation effect size by varying $\sigma_\alpha$ and $\sigma_\beta$.

### S.3 Results from individual-based simulations

We run the IBM for 50,000 timesteps with varying initial values for host discrimination ability $\alpha$ and partner effect $\beta$. The trajectories of the mean trait values in both populations are shown in Figure S1a. For each initial condition, the mean over 10 repetitions was taken to increase robustness of results in the presence of stochasticity. In general, the results corroborate the predictions from the analytical model (see Fig. 2c in the main text), meaning that the trajectories converge to the same two evolutionary attractors, with oscillating behaviour in the neighbourhood of the antagonistic fixed point. Three types of evolutionary transitions can be observed as described in section 3.2 in the main text. Also, varying the cost parameters or relative speed of evolution yielded qualitatively similar results as in the analytical model (not shown).

However, the exact location of the attractors as well as the shape of the basins of attraction slightly deviates from the situation there. For example, trajectories starting with low discrimination ability, but strongly antagonistic partners do not converge to mutualism as often as in the analytical model (see Fig. S1a). This and other quantitative differences are probably effects of intraspecific variation which emphasises the selection also on the edges of the trait distribution in contrast to the deterministic approximation with local selection gradients where selection is considered only infinitesimally close to the mean trait value. Intraspecific trait variation was present but not substantial in the simulations. The standard deviation in the distribution of partner effect $\beta$ was smaller than 0.22 in 99% of all the simulated time steps (calculated at each time step during all simulations described above; 0.26 for host discrimination ability $\alpha$). In comparison, the distance between the two attractors in the trait space is 2.40.

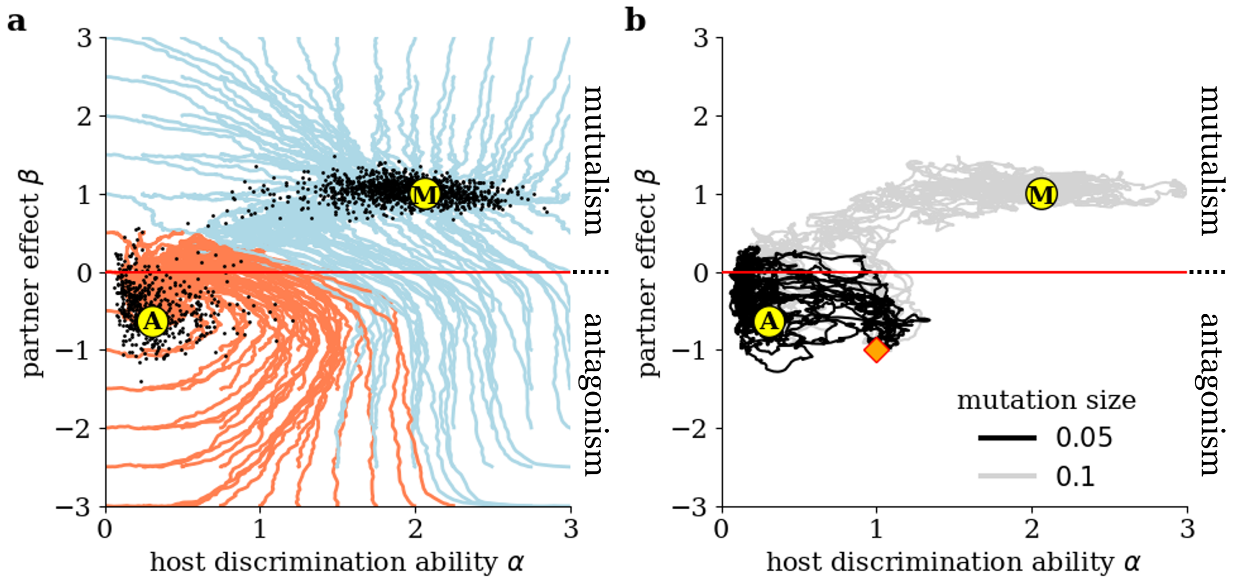

Figure S1: **Coevolutionary trajectories from the individual-based simulations.** (**a**) Trajectories in the trait space for varying initial conditions. Trajectories are averaged over 10 repetitions with the same initial values. Black dots mark the endpoints of single simulation runs. Trajectories are coloured light blue if they end up at the mutualistic attractor (M) and red if they end up at the antagonist one (A). Parameters were chosen according to Table 1 from the main text, with $\sigma_\alpha = \sigma_\beta = 0.05$. (**b**) Trajectories of single simulation runs starting at $\alpha = 1$ and $\beta = -1$ (orange square) for varying mutation size $\sigma_\alpha = \sigma_\beta$. In both panels, for all trajectories a moving average is taken over 200 timesteps for simplicity of visualisation. Yellow circles mark the antagonist and mutualist attractor identified in the analytical model.

In contrast to predictions from the analytical model, if mutation size is increased to $\sigma_\alpha = \sigma_\beta = 0.1$, trajectories tend to ”jump” from the basin of attraction of the antagonistic fixed point to the basin of attraction of the mutualist one. This is visualised in Figure S1b for the 10 repetitions starting with $\alpha = 1$ and $\beta = -1$. For these initial conditions, the analytical model predicts convergence towards the antagonistic attractor. With high mutation sizes, several of the trajectories (in gray) approach the antagonistic fixed point at first but leave its presumed basin of attraction at some point, finally ending up close to the mutualistic fixed point. These jumps do not occur if mutation size is kept at $\sigma_\alpha = \sigma_\beta = 0.05$ (see trajectories in black). Mutation effect size is directly linked to intraspecific trait variation, we therefore deduce that the likelihood of these jumps increases with increasing trait variation. Using the baseline parameterisation, the jumps only happen in the direction from antagonism to mutualism. However, when increasing the cost parameters, jumps in both directions potentially occur (not shown).
